# Differential receptor activation defines the fundamental limit of chemotactic sensing

**DOI:** 10.64898/2025.12.17.694457

**Authors:** Adam Dowdell, David Jordan, Kimia Witte

## Abstract

Cells are often assumed to require a ∼1% difference in ligand concentration across their length to bias chemotaxis, but this Δ*C* framework overlooks the non-linear relation between ligand concentration and receptor activation and, for linear gradients, is uninformative with respect to gradient steepness. Using mathematical modelling and experiments in linear gradients of the non-degradable cAMP analogue Sp-cAMPS, we show that ligand concentration alone does not describe chemotactic behaviour. By expressing signals as the differential proportion of activated receptors (ΔΩ), we find that cells can detect differences as small as 0.026% across their length. Importantly, the minimum differential required to induce bias increases with overall receptor activation, demonstrating that chemotactic sensitivity depends on the receptor occupancy regime rather than on Δ*C*. The true minimal chemotactic threshold therefore lies at concentrations that minimise receptor activation while still eliciting a bias, and is likely to be below 0.026%.

## 2 Introduction

Chemotaxis underpins essential biological processes, including nutrient sensing in unicellular organisms [1], embryonic development [2], and immune cell trafficking [3]. It is also implicated in disease, where excessive leukocyte recruitment contributes to chronic obstructive pulmonary disease and multiple sclerosis [4, 5], and where directed migration of metastatic cells drives the formation of secondary tumours [6], the principal cause of cancer-related mortality [7].

A general framework capable of explaining and ultimately controlling chemotactic behaviour has long been sought. A prevailing model proposes that an ∼ 1% difference in ligand concentration across the cell length (Δ*C*_%_) defines the minimal detectable chemotactic signal [8], a value repeatedly reported in specific experimental contexts [9, 10, 11]. However this threshold applies only under optimised conditions. A fundamental question therefore remains: how do cells actually perceive chemotactic signals, and why do Δ*C*_%_-based measurements fail outside finely optimised gradient regimes.

Cells sense chemotactic cues through membrane-embedded receptors that convert ligand binding into intracellular responses [12]. The proportion – and, crucially, the spatial distribution – of activated receptors depends on ligand concentration, yet this relationship is non-linear except at very low concentrations [13]. A ligand-concentration model is therefore insufficient, as it cannot capture the underlying receptor dynamics that mediate sensing.

Here, we instead develop a receptor-based model, combining mathematical predictions with experimental validation. This builds on our previous work, which showed that active-receptor analysis can account for reverse chemotaxis, directional inversion within gradients, and chemorepulsion arising from competing attractants [14]. We now focus on the critical question of minimal chemotactic sensitivity: the threshold differential in receptor activation across the cell (ΔΩ_*min*_) that is sufficient to generate a directional bias.

## 3 Results

We used a well-established, experimentally tractable chemotaxis system to study the behaviour of cells in a defined linear ligand gradient. When a large source diffuses towards a large sink via a narrow connecting channel, the concentrations in the reservoirs remain effectively unchanged, and a stable gradient forms. This configuration, used in the Insall chemotaxis chamber [15], yields an approximately linear equilibrium gradient in the absence of other influences, providing a controlled system to investigate the threshold parameter ΔΩ_*min*_.

We used aggregation-competent *Dictyostelium discoideum* exposed to the phosphodiesterase-resistant cAMP analogue Sp-cAMPS [16]. A non-degradable ligand is essential to maintain linearity, as degradation by cells would otherwise self-impose exponential gradients [17]. Starved cells express high levels of cAMP receptor 1 (cAR1) peaking at 3-4 hours [18], and this receptor mediates chemotactic signalling at this stage, with ∼7x10^4^ copies per cell [19]. Other cAMP receptors (cAR2, cAR3) are minimally expressed during early starvation, ensuring that cAR1 dominates sensing.

The cAR1 affinity for cAMP, expressed as the concentration at which half of the receptors are bound (*K*_*D*_), is central to translating ligand concentration into receptor activation, but published estimates vary widely, and Sp-cAMPS is less well characterised. Reported values suggest that the *K*_*D*_ for Sp-cAMPS is ∼68-100-fold higher than that of cAMP [20, 21]. Receptor phosphorylation further shifts cAR1 from high cAMP affinity (3.5 - 30*nM* ) to a low-affinity (estimated 200 - 500*nM* ) state, with an approximate 1:9 distribution between them [22, 23, 24]. Based on these constraints, we adopted an Sp-cAMPS *K*_*D*_ of 1*µM* for modelling. The active-receptor (AR) framework can be recalculated for any *K*_*D*_, so this assumption does not limit the generality of the analysis.

### 3.1 Receptor activation defines chemotaxis, not concentration

We performed chemotaxis assays using source Sp-cAMPS concentrations of 1*µM* (low) and 100*µM* (high), each generating a linear gradient to a zero-concentration sink (Fig 1A). Although these gradients differ in steepness by two orders of magnitude, the percentage difference in ligand concentration across a 10*µm* cell (Δ*C*_%_) is identical at all positions along the bridge (Fig 1B). Under a Δ*C*_%_ framework, cells in the 1*µM* and 100*µM* gradients should therefore behave identically. This highlights that Δ*C*_%_ is insensitive to gradient steepness, despite steepness being a key determinant of chemotactic behaviour. Full mathematical details are provided in the supplementary information.

**Figure 1.**
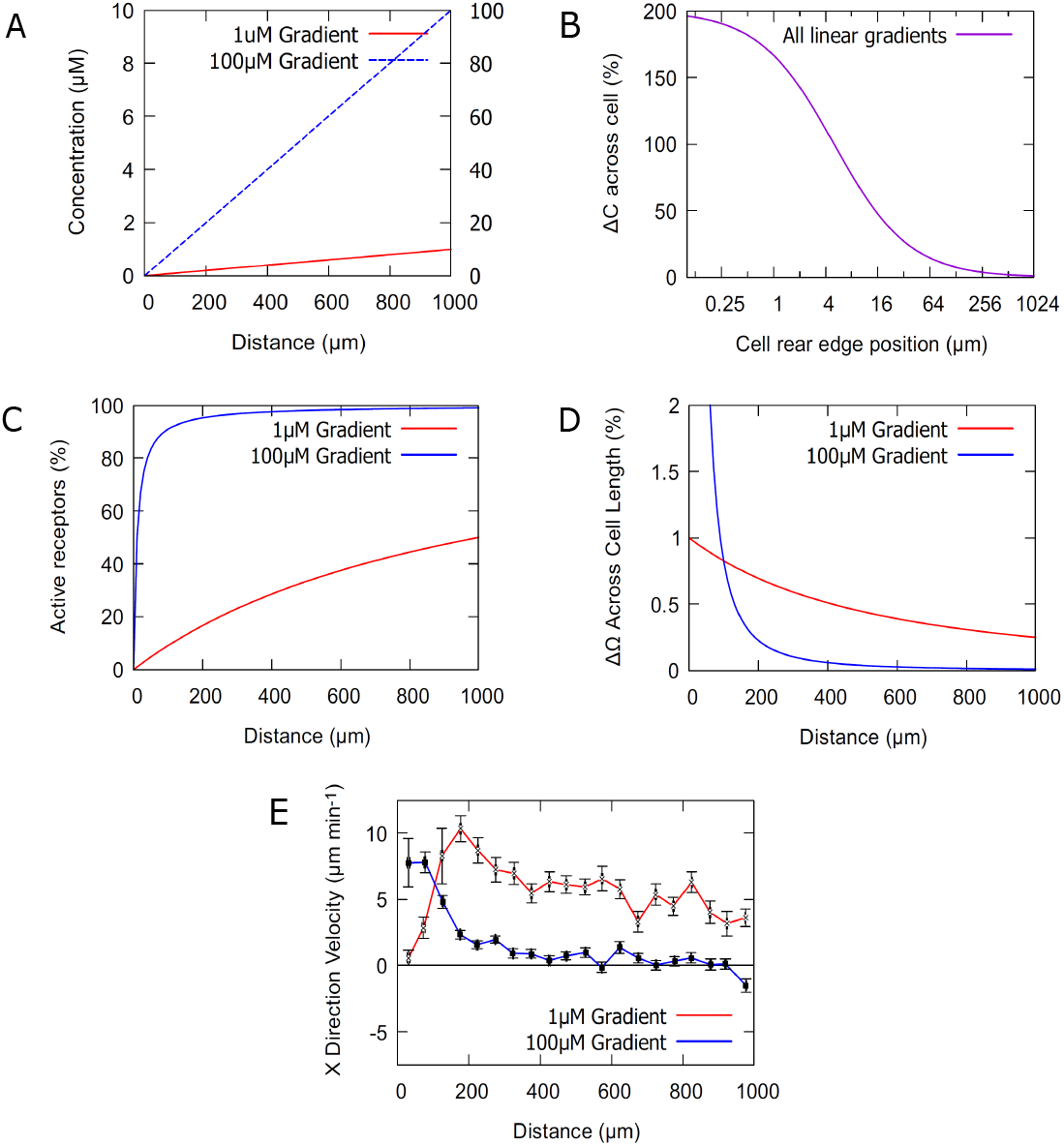
Differential receptor activation, not Δ*C*_%_, distinguishes chemotactic cues. (A) Linear gradients generated from 1*µM* (left-hand y-axis) and 100*µM* (right-hand y-axis) sources, with a bridge length of 1*mm*. (B) Despite differing in steepness by two orders of magnitude, the percentage concentration difference across a 10*µm* cell (Δ*C*_%_) is identical in both gradients, and in all linear gradients. Thus Δ*C*_%_ carries no predictive information about chemotactic behaviour. (C) Active receptor fraction (Ω) along each gradient, predicted by the AR model, showing distinct activation profiles. (D) Differential receptor activation across the cell (ΔΩ) derived from Ω(*x*), for a 10*µm* cell; each gradient produces a unique ΔΩ profile. (E) Experimental chemotaxis data validate AR-model predictions: strong initial bias in the 100*µM* gradient that rapidly declines, and sustained bias across most of the 1*µM* gradient.

Fig 1C shows the proportion of active receptors (Ω) along each gradient. The two profiles are markedly different: in the 100*µM* gradient, Ω reaches 50% very close to the start of the bridge, whereas in the 1*µM* gradient this occurs only at the end, as expected for the chosen *K*_*D*_. The resulting differential in receptor activation across the cell (ΔΩ; Fig 1D) is initially high in the steep gradient, predicting strong chemotactic bias, but collapses rapidly as ligand concentration rises well above *K*_*D*_, falling to ∼0.05% by 40% of the bridge. In contrast, the shallow gradient maintains a substantial and gradually varying ΔΩ of 1-0.25% across the entire bridge. Thus, each gradient generates a distinct Ω(*x*) profile that can be modelled and predicts chemotactic behaviour.

Experimental data (Fig 1E) support these predictions: cells in the 100*µM* gradient show strong initial bias that declines quickly, whereas cells in the 1*µM* gradient retain robust directional migration over ∼90% of the bridge length.

### 3.2 Deriving the minimal chemotactic threshold, ΔΩ_*min*_

To determine the threshold parameter ΔΩ_*min*_, we measured the position at which cells in different gradients no longer displayed a chemotactic bias (Fig 2A, 2B). We defined *x*_*c*_ as the point at which velocity could not be distinguished from zero, i.e. where the lower confidence bound intersected the x-axis. These values were obtained by bootstrap analysis, as described in the supplementary information.

**Figure 2.**
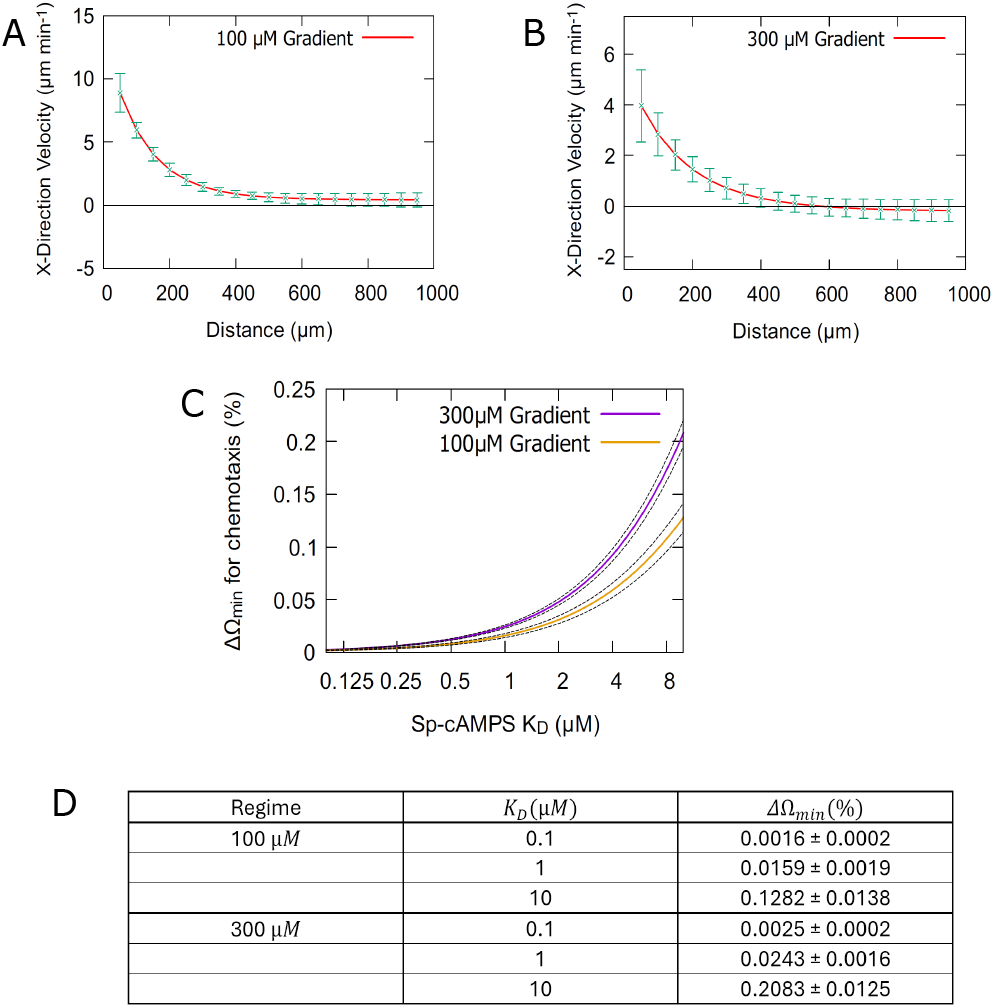
Deriving the minimal detectable receptor-activation differential, ΔΩ_*min*_. (A,B) Bootstrapped velocity profiles (mean ± 95% confidence interval) for cells in 100*µM* and 300*µM* gradients, used to identify the position *x*_*c*_ at which chemotaxis becomes indistinguishable from zero. (C) Relationship between Sp-cAMPS *K*_*D*_ and ΔΩ_*min*_, obtained by applying the AR model to *x*_*c*_ values from both gradients, based on a minimum of three experimental repeats. (C) Inferred ΔΩ_*min*_ values across a range of modelled *K*_*D*_ values, showing systematic dependence on receptor affinity.

Using these *x*_*c*_ values, we applied the AR framework to derive a general relationship between *K*_*D*_ and ΔΩ_*min*_ (Fig 2C). Across all *K*_*D*_ values tested, ΔΩ_*min*_ was larger for the 300*µM* gradient, demonstrating that the minimum differential required to induce a bias increases with ligand concentration and overall receptor activation. Uncertainty in the precise *K*_*D*_ of Sp-cAMPS does not limit the analysis: ΔΩ_*min*_ can be computed for *K*_*D*_ values from 0.1 - 10*µM* (and beyond), and varies systematically with receptor affinity (Fig 2D). This behaviour is expected, as changing *K*_*D*_ alters total receptor occupancy and therefore the percentage difference between the cell front and the rear. Assuming a *K*_*D*_ of 1*µM*, these analyses place an upper bound on ΔΩ_*min*_ of approximately 0.026%.

### 3.3 Mapping chemotaxis across arbitrary chemical gradients

In steep gradients, cells cease migrating when they can no longer resolve the signal, i.e. when ΔΩ across the cell falls below the minimal detectable level. Fig 2D shows that, assuming a *K*_*D*_ of 1*µM*, the upper bound on this signal is ΔΩ_*min*_ ≈0.026%. Thus, locations where ΔΩ *>* ΔΩ_*min*_ should support chemotaxis. We therefore generated predictive heatmaps of ΔΩ, and the corresponding behavioural regime at every point along a broad range of linear gradients (Fig 3A, 3B). In Fig 3A, gradients in the range 1 - 1000*µM* are plotted. Magenta indicates regions where ΔΩ *<* ΔΩ_*min*_ and chemotaxis fails due to receptor saturation. Blue and cyan denote regions where ΔΩ *>* ΔΩ_*min*_, supporting weaker chemotactic regimes of increasing strength. Yellow marks regions ΔΩ *>>* ΔΩ_*min*_, where chemotaxis is strong. The black band identifies positions in each gradient where ΔΩ = ΔΩ_*min*_.

**Figure 3.**
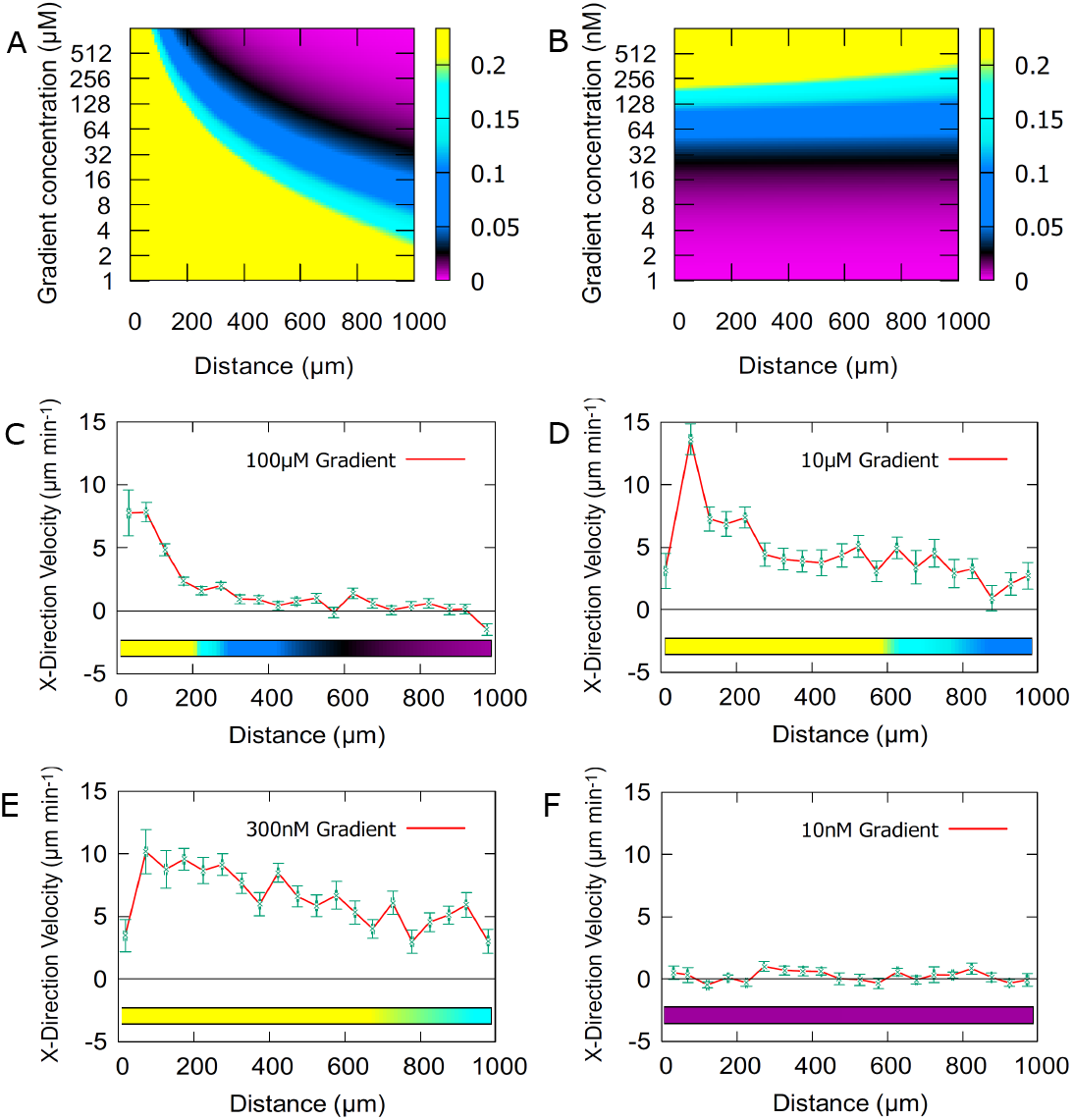
Predictive chemotactic regimes with ligand concentration, validated experimentally. (A) Predicted ΔΩ and chemotactic behaviour along gradients ranging 1 - 1000*µM* (*K*_*D*_ = 1*µM* ). Magenta indicates ΔΩ *<* ΔΩ_*min*_ and loss of chemotaxis due to receptor saturation; blue and cyan denote ΔΩ *>* ΔΩ_*min*_ and weaker chemotaxis; yellow marks strong chemotaxis where ΔΩ *>>* ΔΩ_*min*_. The black band shows positions where ΔΩ = ΔΩ_*min*_. (B) Prediction for gradients spanning 1 - 1000*nM*, where chemotaxis fails (magenta) due to insufficient receptor activation in very shallow gradients. (C-F) Experimental migration profiles across gradients from 10*nM* to 100*µM* confirm model predictions, with behavioural regimes matching the corresponding ΔΩ-based classifications.

Fig 3B shows predicted cell behaviour at concentrations from *K*_*D*_ down to 1000-fold lower. In this regime, the magenta zone again denotes failure of chemotaxis, now due to insufficient receptor activation in extremely shallow gradients.

We tested these predictions experimentally across gradients spanning 10*nM* to 100*µM* (Fig 3C-F). The observed behaviours agree well with the model, as indicated by the coloured bars beneath each curve. Notably, both the 300*nM* and 10*µM* gradients support strong chemotaxis across the full bridge, with the 10*µM* gradient weakening only at its upper end. This pattern is consistent with the true Sp-cAMPS *K*_*D*_ lying between these concentrations. A small positive bias is observed in the region of ΔΩ = 0.026% in the 100*µM* gradient, consistent with expectation: ΔΩ_*min*_ decreases at lower overall receptor activation, and the value ΔΩ = 0.026% derives from a 300*µM* gradient.

## 4 Discussion

Examination of the mathematical definition of Δ*C*_%_ reveals clear limitations in ligand concentration-based models of chemotactic sensitivity. This framework has persisted because it approximates behaviour under highly optimised chemical conditions, but outside such regimes there is no consistent relationship between Δ*C*_%_ and chemotactic response. In contrast, the active-receptor (AR) model accurately describes and predicts chemotaxis across many orders of ligand conditions, provided that *K*_*D*_ is at least approximately known. Precise knowledge of *K*_*D*_ is unnecessary, and *K*_*D*_ itself can be inferred by comparing predicted and observed chemotactic dynamics.

The central strength of the AR model is that, despite its powerful ability to incorporate multiple ligands and gradients, its output reduces to a single organising parameter: ΔΩ. The strength and direction of the response are then simply the magnitude and sign of ΔΩ, respectively. Essentially, this parameter governs all key features of chemotactic behaviour, irrespective of how many ligands contribute to that signal.

Most chemotaxis assays use ligand concentrations that do not saturate receptors. Instead employing gradients that saturate at different positions along the bridge (Fig 2), we identify for each a gradient position *x*_*c*_ at which chemotaxis ceases. When *K*_*D*_ is known, *x*_*c*_ yields the minimal detectable receptor-activation differential, ΔΩ_*min*_; when *K*_*D*_ is uncertain, the relationship between *K*_*D*_ and ΔΩ_*min*_ can still be derived. This analysis reveals a striking result: ΔΩ_*min*_ increases in higher gradients, meaning that greater receptor-activation differences are required for bias when overall receptor occupancy is high. In saturated environments, therefore, cells become less sensitive to directional information.

Assuming a *K*_*D*_ of 1*µM*, we estimate ΔΩ_*min*_ = 0.024% for 300*µM* gradients and 0.016% for 100*µM* gradients. Because determining ΔΩ_*min*_ requires an experimentally measurable *x*_*c*_, it cannot be directly obtained for lower, non-saturating concentrations. However, the monotonic decrease observed between 300*µM* and 100*µM* suggests that ΔΩ_*min*_ would fall for sub-saturating concentrations, likely below 0.016%. A fundamental lower bound must exist: at sufficiently low occupancy, a difference of one active receptor across the cell defines the resolvable ΔΩ. Below this limit, no further increase in sensitivity is possible. Thus, cells cannot become arbitrarily sensitive merely by reducing overall receptor activation.

As shown by the heatmap predictions and experimental validation (Fig 3), the AR framework remains accurate across at least 4 orders of magnitude in ligand concentration. At very high concentrations, many hundreds of times the *K*_*D*_, receptors saturate almost immediately, and the chemotactic window becomes extremely narrow; in this regime, little bias is expected, regardless of gradient shape. Fig 3C shows that cells can just detect ΔΩ = 0.026% when *K*_*D*_ = 1*µM* . If *K*_*D*_ is instead 10*µM*, the same ΔΩ corresponds to gradients of the order 300*nM* rather than 100*µM* (as illustrated in the supplementary information), with Fig 3E showing clear bias. For *K*_*D*_ = 50*µM* even smaller signals are elicited in the same gradient regime. This agreement for *K*_*D*_ ≥ 1*µM* supports the conclusion that cells can detect receptor-activation differences as small as 0.026%. Assuming there are of the order 1x10^4^ receptors located at the cell periphery, these signals could represent differentials as low as 2 or 3 receptors across the cell length.

The AR model is readily applied to different experimental configurations, independent of ligand or cell type, and does not require precise *K*_*D*_ values to generate reliable predictions. It therefore provides a general, quantitative framework for modelling chemotaxis.

## Acknowledgements

Supported by the ARIA “Nature Computes Better” Opportunity Space, (Bio)active Matter Based Computation. K.W. conceived the project, supervised and obtained funding. A.D. originated the active-receptor framework for chemotaxis and signalling-limit concept. A.D led experiments and data analysis. A.D. wrote the original draft and D.J. and K.W. revised the manuscript.

