## Supplementary Information for "Differential receptor activation defines the fundamental limit of chemotactic sensing"

### SI–Theory: Mathematical Framework

#### Figure 1

Diffusion from regions of high to low concentration is described by

$$\frac{\partial C}{\partial t} = D\nabla^2 C, \tag{1}$$

where $C$ is concentration, $t$ is time, $D$ is the diffusion coefficient and $\nabla^2$ is the Laplacian. This can be simplified by assuming transport is effectively one-dimensional along $x$, from source to sink.

$$\nabla^2 = \frac{\partial^2 C}{\partial x^2}, \tag{2}$$

so equation (1) becomes

$$\frac{\partial C}{\partial t} = D\frac{\partial^2 C}{\partial x^2}. \tag{3}$$

At steady state, the left-hand side of equation (3) is zero, giving

$$\frac{\partial^2 C}{\partial x^2} = 0, \tag{4}$$

which is solved by integrating twice with respect to $x$ to obtain

$$C(x) = Ax + B, \tag{5}$$

with constants of integration $A$ and $B$. The constants are determined by boundary conditions. The sink at $x = 0$ is fixed at zero concentration and the source at $x = L$ at concentration $C_f$:

$$C(0,t) = 0 \quad , \quad C(L,t) = C_f. \tag{6}$$

Applying these conditions yields:

$$B = 0 \quad , \quad A = \frac{C_f}{L}, \tag{7}$$

and substituting into equation (5) gives the steady-state gradient

$$C(x) = \frac{C_f x}{L}. \tag{8}$$

This profile is linear.

#### Panel B

For the linear gradient in equation (8), the concentration difference between points $x_2$ and $x_1$ ($x_2 > x_1$) is

$$\Delta C = \frac{C_f x_2}{L} - \frac{C_f x_1}{L} = \frac{C_f}{L}(x_2 - x_1). \tag{9}$$

If $x_2$ and $x_1$ correspond to the front and back of a cell, define the cell width $W = x_2 - x_1$, giving

$$\Delta C = \frac{C_f W}{L}. \tag{10}$$

For a linear gradient, the mean concentration at the cell location, $C_{av}$, is the average of the concentrations at the cell extrema:

$$C_{av} = \frac{1}{2}\left(\frac{C_f x_2}{L} + \frac{C_f x_1}{L}\right) = \frac{C_f}{2L}\left(x_2 + x_1\right), \tag{11}$$

Using $x_2 = W + x_1$, equation (11) can be rewritten as

$$C_{av} = \frac{C_f}{2L}(W + 2x_1). \tag{12}$$

The percentage difference across the cell, $\Delta C_{\%}$, relative to the mean concentration is

$$\Delta C_{\%} = \frac{\Delta C}{C_{av}} * 100 = \frac{200W}{W + 2x_1} \tag{13}$$

This expression is independent of both final concentration and gradient length; thus $\Delta C_{\%}$ cannot account for chemotactic sensing and motivates an alternative model.

#### Panel C

To relate an arbitrary set of ligand concentrations ($n$ ligands) to receptor binding, consider unbound receptors $U$ binding ligands $C_i$ to form complexes $G_i$ for $i \in [1, n]$:

$$U + C_i \ \ k_i \longleftrightarrow k_{-i} \ \ G_i. \tag{14}$$

Here, $k_i$ and $k_{-i}$ are the association and dissociation rates for ligand $i$. By the law of mass action, reaction rates scale with the product of reactant concentrations, giving for $U$:

$$\frac{\partial U}{\partial t} = -k_i U C_i + k_{-i} G_i. \tag{15}$$

Analogous expressions follow for $C_i$ and $G_i$. Assuming equilibrium, binding and dissociation balance and $\frac{\partial U}{\partial t} = 0$; equivalently, the free receptor pool is time-independent. This assumption simplifies the analysis and is justified by experimental validation of the resulting predictions. Substituting $\frac{\partial U}{\partial t} = 0$ into equation (15) gives

$$\frac{UC_i}{G_i} = \frac{k_{-i}}{k_i} = K_{D_i}. \tag{16}$$

$K_{D_i}$ is the dissociation constant for ligand $i$, i.e. the ligand concentration at which half the receptors are bound. Setting $C_i = K_{D_i}$ yields $U = G_i$, implying equal free and bound receptor counts and therefore 50% occupancy. Let $\alpha_i \in [0, 1]$ denote the intrinsic efficacy (proportion of receptors activated upon binding) of ligand $i$. For n ligands competing for the same receptor, overall receptor activation must reflect active, not merely bound, receptors. Define $\Omega$ as the efficacy-weighted fraction of receptors that are bound:

$$\Omega = \frac{\sum_{i=1}^{n} \alpha_i G_i}{U + \sum_{i=1}^{n} G_i}. \tag{17}$$

Substituting $G_i$ using equation (16) gives

$$\Omega(C) = \frac{\sum_{i=1}^{n} \frac{\alpha_i C_i}{K_{Di}}}{1 + \sum_{i=1}^{n} \frac{C_i}{K_{Di}}}. \tag{18}$$

Thus, $\Omega = 0.7$ indicates a ligand mixture that yields 70% active receptors. If only Sp-cAMPS is present, a full agonist with $\alpha = 1$, then receptor activation at concentration $C$ is

$$\Omega(C) = \frac{\frac{C}{K_D}}{1 + \frac{C}{K_D}} = \frac{C}{C + K_D}. \tag{19}$$

Here $K_D$ is the Sp-cAMPS dissociation constant and $C$ is the local Sp-cAMPS concentration. Because Sp-cAMPS is engineered to be non-degradable, the gradient approaches a linear steady state, so

$$C(x) = \frac{C_f x}{L}, \tag{20}$$

where $L$ is gradient length and $C_f$ is the final concentration. Substituting equation (20) into equation (19) yields receptor activation as a function of position:

$$\Omega(x) = \frac{\frac{C_f x}{L}}{K_D + \frac{C_f x}{L}} = \frac{C_f x}{K_D L + C_f x}. \tag{21}$$

For an assumed Sp-cAMPS $K_D$, this expression can be evaluated across $x$ for different final concentrations $C_f$. As a percentage,

$$\Omega_{\%} = \frac{C_f x}{K_D L + C_f x} * 100. \tag{22}$$

### Panel D

To convert $\Omega(x)$ into the across-cell difference in active receptors at each position, $\Delta\Omega$, differentiate equation (21) with respect to $x$:

$$\frac{\partial \Omega}{\partial x} = \frac{C_f K_D L}{(K_D L + C_f x)^2}. \tag{23}$$

At position $x$, the derivative relates to $\Delta\Omega$ over cell length $W$ via

$$\frac{\partial \Omega}{\partial x} = \frac{\Delta\Omega}{W}. \tag{24}$$

Substituting equation (24) into equation (23) gives

$$\Delta\Omega = \frac{W C_f K_D L}{(K_D L + C_f x)^2}. \tag{25}$$

In percentage form,

$$\Delta\Omega_{\%} = \frac{W C_f K_D L}{(K_D L + C_f x)^2} * 100, \tag{26}$$

This can be evaluated across $x$ for any specified gradient.

#### Figure 2

##### Panels A, B

To estimate where receptor-saturating conditions lead to cessation of chemotaxis, a statistically rigorous criterion was required. Identifying where the mean velocity first reaches zero ignores variability in neighbouring data, which often oscillates around the x-axis (see main text Fig 3C). Accordingly, a bootstrap analysis was used to leverage a broader data range when estimating the first location of zero cell velocity. Spatially partitioned velocity data were resampled with replacement 1000 times, and each resample was fit with

$$f(x) = Ae^{-bx} + C. \tag{27}$$

Often, one or more points near the beginning and end of the gradient deviate from expected behaviour (main text Fig 3A, 3B). These points frequently show elevated uncertainty, indicating altered or unstable responses at the gradient extrema. A detailed investigation of this behaviour is beyond the scope of this work; it may reflect transient chemical or spatial adaptation as cells enter or exit the gradient, or arise from Insall chamber geometry. Regardless, to estimate where chemotaxis ceases, edge data are excluded so that fits are applied only to data that conform to the model and its expected form. Fit parameters $A$, $b$ and $C$ were recorded for each resample. Each fitted function was then evaluated at $50\mu m$ increments along the gradient (excluding the edge regions) to generate velocity values. Across 1000 resamples, these values were aggregated to form an overall mean fit and a 95% confidence interval (CI). Chemotaxis cessation was defined as the position where the lower-bound CI intersects the $x$-axis, i.e. where zero velocity cannot be statistically excluded. To obtain $x_c$, the lower-bound CI was fit with the same functional form as equation (27), and $x_c$ was solved via $f(x) = 0$. This procedure was applied to each experimental repeat for the $100\mu M$ and $300\mu M$ conditions, and the mean $x_c$ and standard error on the mean (SEM) were computed (Table 1).

| | 300 μ*M* Stop Point ($x_c$) | 100 μ*M* Stop Point ($x_c$) |
|---|---|---|
| | 389.0 | 670.6 |
| | 381.3 | 815.9 |
| | 369.6 | 864.0 |
| | 326.9 | |
| Mean | 366.7 | 783.5 |
| Standard deviation | 24.0 | 82.21 |
| SEM | 12.0 | 47.5 |
| Mean ± SEM | 366.7 ± 12.0 | 783.5 ± 47.5 |

Table 1: Estimated cessation position $x_c$ for all repeats under $100\mu M$ and $300\mu M$ conditions, including standard deviation and SEM.

#### Panel C

If chemotaxis ceases at a critical position $x_c$, then the across-cell difference in active receptors at that position corresponds to the minimum directional cue, $\Delta\Omega_{min}$. If $x_c$ can be experimentally determined, then

$$\Delta\Omega_{min}(\%) = \frac{WC_f K_D L}{(K_D L + C_f x_c)^2} * 100. \tag{28}$$

At this stage, all parameters are known except the Sp-cAMPS $K_D$ and the minimum signal $\Delta\Omega_{min}$, so a relationship can be drawn between them. Once $K_D$ is determined, it maps to a specific $\Delta\Omega_{min}$. Across repeated experiments at a fixed final concentration $C_f$, a distribution of $x_c$ values is obtained, allowing estimation of the mean $x_c$ and its SEM, $\sigma_{x_c}$ (Table 1). An uncertainty in $x_c$ propagates to $\Delta\Omega_{min}$ via the $x$-dependence of equation (25). Differentiating equation (25) with respect to $x$ gives

$$\frac{\partial \Delta\Omega}{\partial x} = \frac{-2WC_f^2 K_D L}{(K_D L + C_f x)^3}. \tag{29}$$

Because this derivative quantifies how $\Delta\Omega$ varies with $x$, $\sigma_{\Delta\Omega_{min}}$ scales with $\sigma_{x_c}$ accordingly. Therefore,

$$\sigma_{\Delta\Omega_{min}}(\%) = \left| \frac{-2WC_f^2 K_D L}{(K_D L + C_f x)^3} \right| \sigma_{x_c} * 100. \tag{30}$$

Uncertainty bands for curves generated from equation (28) are obtained by plotting

$$\Delta\Omega_{min}(\%) \pm \sigma_{\Delta\Omega_{min}}(\%) = \left( \frac{WC_f K_D L}{(K_D L + C_f x_c)^2} \pm \frac{2WC_f^2 K_D L}{(K_D L + C_f x)^3} \sigma_{x_c} \right) * 100. \tag{31}$$

### Figure 3

#### Panels A, B

For a specified $K_D$, $\Delta\Omega_{min}$ can be inferred from equation (28) using the cessation coordinate $x_c$. A heat map can then be generated to predict how chemotactic cues vary across concentration regimes by plotting equation (26) in three dimensions: position $x$ on the $x$-axis, final concentration $C_f$ on the y-axis (over the desired orders of magnitude), and $\Delta\Omega_{\%}$ on the $z$-axis. Rendering was applied to $z$-axis values using the defined $\Delta\Omega_{min}$ range to indicate whether chemotaxis is expected at a given position under a given gradient condition.

#### Discussion

To support the hypothesis that cells can detect $\Delta\Omega = 0.026\%$, an additional graph is provided for a $300nM$ gradient assuming Sp-cAMPS $K_D$ values of $10\mu M$ and $50\mu M$. Comparison of Fig 1 with Fig 3E (main text) indicates that signals of this magnitude can still induce a chemotactic bias when $K_D = 10\mu M$. Increasing to $K_D = 50\mu M$, even smaller signals are elicited in this same gradient, with clear bias. Overall, these results indicate that $\Delta\Omega = 0.026\%$ is sufficient to provide directional cues for Sp-cAMPS $K_D \geq 1\mu M$.

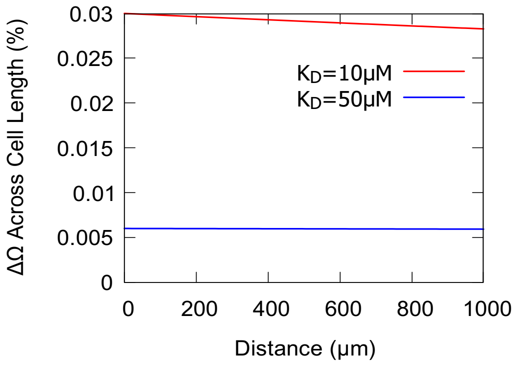


Figure 1: $\Delta\Omega_{\%}$ across a $300nM$ gradient assuming $K_D$ values of $10\mu M$ and $50\mu M$.

### SI–Methods: Experimental and Analytical Methods

#### Microorganisms

- *Klebsiella aerogenes.*
- Adenyl cyclase knockout *Dictyostelium discoideum* ($ACA^-$).

#### Reagents

**KK2 buffer.** A 10× stock was prepared by dissolving 22 g $KH_2PO_4$ (monobasic) and 7 g $K_2HPO_4$ (dibasic) in 1 L distilled $H_2O$, autoclaved for long-term storage, and diluted to 1× for use.

**$CaCl_2$/$MgCl_2$ salts.** A 100× stock was prepared by dissolving 2.22 g $CaCl_2$ and 3.81 g $MgCl_2$ in 200 mL distilled $H_2O$, autoclaved, and added to KK2 to 1× where indicated (KK2 + salts denoted KK2MC).

**SM agar plates.** SM agar premix was prepared according to the manufacturer's instructions, autoclaved, and poured thickly (∼35 mL per 90 mm plate). Plates were used for growth and passaging of both microorganisms.

**LB broth.** Lennox-formula LB broth was used to prepare bacterial suspensions and to spread bacterial lawns on SM agar plates (∼500 $\mu$L per 90 mm plate), followed by brief drying.

#### Passaging

***K. aerogenes.*** Bacteria were passaged weekly by streaking onto fresh SM agar plates. Stocks were replaced monthly.

***D. discoideum* ($ACA^-$).** $ACA^-$ cells were passaged once or twice weekly by inoculating a fresh bacterial lawn using vegetative cells from the feeding front. Fruiting bodies do not form in $ACA^-$ mutants due to inability to produce cAMP; plates lacking a clear feeding front were discarded. Stocks were replaced monthly.

#### Experiment preparation

*D. discoideum* cultures were prepared two days in advance by generating four bacterial suspensions containing vegetative $ACA^-$ cells, each diluted four-fold relative to the previous, and spreading each suspension onto a separate SM plate.

#### Pulsed chemotaxis assay

$ACA^-$ cells were harvested from plates displaying a uniform "glassy" appearance (indicative of bacterial consumption and eukaryote proliferation). Plates or regions with a textured appearance were excluded. Cells were harvested by adding 5 mL KK2 to the plate, scraping, and transferring the suspension to a container; KK2 was added to a final volume of 20 mL.

Cells were washed three times by centrifugation at 300 g for 3 min, discarding supernatant and resuspending in 20 mL KK2 between washes. The final suspension was vortexed, counted, and adjusted to $1 \times 10^7$ cells $mL^{-1}$ in KK2. A total of 20 mL of this suspension ($2 \times 10^8$ cells) was placed in a beaker, covered with foil to limit evaporation while allowing gas exchange, and gently shaken at 120 rpm (or gently stirred) for 1 h. In parallel, 20 mL of 20 $\mu$M cAMP in KK2 was prepared.

After 1 h, cells were pulsed with 100 nM cAMP every 6 min for 4 h using a programmable dual-syringe pump (while still shaking/stirring). Aliquots (1 mL) were taken for each condition (up to four per batch). Each aliquot was washed three times in an Eppendorf centrifuge using the short-spin function (∼5 s), aspirating supernatant and resuspending in KK2MC each time.

One suspension was vortexed and diluted 1:20 in KK2MC, vortexed again, and plated as 200 $\mu$L onto 22×22 mm glass coverslips (four coverslips), then incubated for 20 min to allow adhesion. Remaining suspensions were stored at 4°C. During adhesion, KK2MC solutions containing the desired Sp-cAMPS concentrations were prepared.

Insall chambers were assembled by filling indentations and channels with KK2MC such that a small meniscus was visible. Excess liquid was removed from the coverslip, which was placed cell-side down over the chamber, leaving the tips uncovered. Liquid contained in the outer channel was aspirated via an exposed tip, and Sp-cAMPS solution was introduced into this outer channel to establish a stable linear gradient between the source (outer channel) and the sink (centre).

Cell density on the bridge was checked and adjusted by modifying dilution where required; chambers were remade if cells were not positioned appropriately. After 10 min equilibration, the bridge region was imaged for ≥20 min at 15–30 s intervals. Additional slides were prepared sequentially from refrigerated suspensions for remaining conditions.

#### Analysis

Cell trajectories were extracted using a custom FIJI plugin (Luke Tweedy). Positions were converted from pixels to $\mu$m, with $x = 0$ defined at the start of the bridge (and the gradient). For each track, mean position and $x$-directed velocity were calculated. Tracks were binned into 50 $\mu$m intervals by mean $x$-position, and per-bin mean velocity and SEM were computed for all velocity analyses.
